# QTLs for potato tuber resistance to *Dickeya solani* are located on chromosomes II and IV

**DOI:** 10.1101/2021.02.19.432067

**Authors:** Renata Lebecka, Jadwiga Śliwka, Anna Grupa-Urbańska, Katarzyna Szajko, Waldemar Marczewski

**Affiliations:** Plant Breeding and Acclimatization Institute, National Research Institute, Młochów Research Center, Platanowa 19, 05-831 Młochów, Poland

**Author notes:** **The e-mail addresses, telephone and fax numbers of corresponding authors**, R. Lebecka, Jadwiga Śliwka.

**Keywords:** potato soft rot resistance, *Dickeya solani* mapping, diploid hybrids of *Solanum* spp., starch

## Abstract

Soft rot is a bacterial disease that causes heavy losses in potato production worldwide. The goal of this study was to identify quantitative trait loci (QTLs) for potato tuber resistance to bacterium *Dickeya solani* and for tuber starch content to study the relationship between these traits. A highly resistant diploid hybrid of potato was crossed with a susceptible hybrid to generate the F1 mapping population. Tubers that were wound-inoculated with bacteria were evaluated for disease severity expressed as the mean weight of rotted tubers, and disease incidence measured as the proportion of rotten tubers. Diversity array technology (DArTseq^™^) was used for genetic map construction and QTLs analysis. The most prominent QTLs for disease severity and incidence were identified in overlapping regions on potato chromosome IV and explained 22.4% and 22.9% of the phenotypic variance, respectively. The second QTL for disease severity was mapped to chromosome II and explained 16.5% of the variance. QTLs for starch content were detected on chromosomes III, V, VI, VII, VIII, IX, XI, and XII in regions different than the QTLs for soft rot resistance. Two strong and reproducible QTLs for resistance to *Dickeya solani* on potato chromosomes IV and II might be useful for further study of candidate genes and marker development in potato breeding programs. The relationship between tuber resistance to bacteria and the starch content in potato tubers was not confirmed by QTL mapping, which makes the selection of genotypes highly resistant to soft rot with a desirable starch content feasible.

## Introduction

Bacteria of the genera *Pectobacterium* and *Dickeya* are broad host range pathogens; together, they cause soft rot of species from 50% of angiosperm plant orders and are among the top 10 most important phytopathogenic bacteria (Ma et al. 2007; Mansfield et al. 2012). These bacteria are ubiquitous in the environment and opportunistic in potato plants (*Solanum tuberosum* L.). *Dickeya* species are either facultative saprophytes or aggressive necrotrophs, and are considered hemibiotrophs that cause severe losses, yield reduction, and pre- and postharvest decay of potato tubers (Toth et al. 2011; Hugouvieux-Cotte-Pattat 2016; Kraepiel et al. 2016). Potato tubers can be infected by bacteria through wounds, natural openings (stomata, lenticels), or stolons. *D. solani* can cause disease of potato even with a low inoculum level, is more likely to colonize potato and is more aggressive than other pectinolytic bacteria (Czajkowski 2009; Toth et al. 2011). The most effective strategies to control bacterial soft rot in potato crops are the application of sanitation measures and enhancement of potato resistance, while chemical control is seldom used for these bacterial diseases. However, there are few highly resistant commercial potato cultivars, and the complexity of the inheritance of resistance to soft rot bacteria makes breeding potato cultivars with high levels of resistance difficult (Bethke et al. 2019; Charkowski et al. 2020). To date, there is no evidence of any species/strain–specific resistance of potato to soft rot bacteria. In addition, there are no examples of gene-for-gene resistance for *Dickeya* spp. and the basis for resistance to *Dickeya* spp. in wild potato species is poorly understood (Charkowski et al. 2020).

Three different sources of high resistance to *P. atrosepticum* with some degree of common origin have been described so far at IHAR-PIB: the 1^st^ source, represented by clone DG 83-2025, consists of *S. tuberosum* (47.7%), *S. chacoense* (43.8%), and *S. yungasense* (8.6%) (Zimnoch-Guzowska et al. 2000); the 2^nd^ source, represented by clone DG 88-9, consists of *S. tuberosum* (47%), *S. phureja* (25.3%), *S. chacoense* (15.8%), *S. acaule* (4.7%), *S. yungasense* (3.1%), *S. gourlayi* (3.1%) and *S. demissum* (1%) (Lebecka et al. 2004), and the 3^rd^ source, represented by clone DG 00-270, consists of *S. tuberosum* (47.3%), *S. phureja* (21.9%), *S. chacoense* (19.9%), *S. yungasense* (5.5%), *S. verrucosum* (3.1%) and *S. microdontum* (2.3%). The clone DG 00-270 is partly related to the 1^st^ source of resistance, diploid DG 83-2025, has the common progenitors *S. chacoense* (12.5%) and *S. yungasense* (5.5%) and is partly related to the 2^nd^ described source of resistance, DG 88-9, which shared common progenitors from *S. phureja* (12.5%), *S. chacoense* (13.3%), and *S. yungasense* (3.5%) (Lebecka et al. 2020). We do not know which species of *Solanum* was the source of high resistance to soft rot bacteria, it may be *S. chacoense*, as it was suggested by Zimnoch-Guzowska and Łojkowska (1993).

In diploid potato, the first QTL mapping experiments for tuber and foliar resistance to *P. atrosepticum* (syn. *Erwinia carotovora* subsp. *atroseptica*) were carried out in hybrids originating from the maternal clone DG 83-2025, representing the 1^st^ source of resistance (Zimnoch-Guzowska et al. 2000). The mapping results revealed the complex polygenic nature of this trait. The high resistance of DG 88-9, representing the 2^nd^ source of resistance, was transferred to tetraploid potato clones (Lebecka and Zimnoch-Guzowska 2004). Subsequently, the tetraploid clone E97-159 was crossed with the cultivar Herta in a commercial breeding program, resulting in the potato cultivar Mieszko, which was registered in 2015 in Poland. This cultivar showed a higher level of partial resistance to *D. solani* than 10 other potato cultivars (Lebecka and Michalak 2017).

The objective of this study was to describe the genetic factors underlying potato tuber resistance to soft rot caused by *D. solani* in the diploid clone DG 00-270 through QTL analyses using DArTseq^™^ markers and linkage map construction.

There are reports that potato cultivars with a high content of dry matter are less susceptible to bacterial soft rot than tubers with lower dry matter contents; however, the relationship between the resistance and tuber starch content is not clear (Biehn et al. 1972; Tzeng et al. 1990; Wright et al. 2005; Zimnoch-Guzowska and Łojkowska 1993). A study in unselected diploid populations of *Solanum* spp. revealed a lack of correlation between tuber soft rot and starch content (Lebecka and Zimnoch-Guzowska 2004), and in contrast, a significant correlation (−0.46) was detected between lesion diameter and dry matter content in a population obtained after crossing the highly soft rot resistant *S. chacoense* Bitter clone M6 with the susceptible *S. tuberosum* dihaploid clone US-W4 (Chung et al. 2017). In the current study, we report the first QTL map for resistance to *D. solani* infection, which was not correlated with tuber starch content.

## Materials and methods

### Plant material

A segregating F1 diploid potato population, referred to as DS-13, consisting of 176 individuals was used for QTL mapping. The population was derived from the cross between two diploid potato clones, DG 00-270 (the resistant maternal parent, P1) and DG 08-305 (the susceptible pollen parent, P2) (Lebecka et al. 2019; Lebecka et al. 2020). The parental clones were bred at the Plant Breeding and Acclimatization Institute-National Research Institute (IHAR-PIB), Młochów Research Center and were complex interspecific hybrids of *Solanum* species. P1 had theoretical genetic backgrounds of *Solanum tuberosum* L. (47.7%), *S. chacoense* Bitter (19.5%), *S. phureja* Juz. and Bukasov (21.9%), *S. yungasense* Hawkes (5.5%), *S. verrucosum* Schltdl (3.1%), and *S. microdontum* Bitter (2.3%), and P2: *S. tuberosum* (63.5%), *S. chacoense* (11.1%), *S. phureja* (14.2%), *S. yungasense* (3.5%), *S. verrucosum* (0.4%), *S. gourlayi* Hawkes (3.5%), *S. acaule* Bitter (1.6%), *S. stenotomum* Juz., and Bukasov (1.6%), *S. demissum* Lindl. (0.5%). The P1 parent was characterized as fertile (50% of pollen grains stained with lacto phenol acid fuchsine) and had the ability to produce unreduced male gametes (large pollen grains) (Lebecka et al. 2020). Two potato cultivars with different levels of resistance to soft rot were included in tests as standards by which to assess resistance to *D. solani*: the table cultivar Irys was highly susceptible, scoring as 2, and the high-starch cultivar Glada was moderately resistant, scoring as 5 on a scale of 1 to 9, where 9 is the most resistant, according to the descriptions of potato cultivars registered in Poland.

### Bacterial inoculum

The IFB0099 strain, syn. IPO2276, (IPO: Plant Research International Collection, Wageningen, The Netherlands) of *D. solani*, was used for inoculation. This strain was isolated in Poland in 2005 from a cultivar Sante plant exhibiting blackleg (Slawiak et al. 2009; Golanowska et al. 2015) and was obtained from the collection of the Intercollegiate Faculty of Biotechnology University of Gdańsk and Medical University of Gdańsk, Poland. After 24 h of incubation on LB agar medium, the bacteria (stored at −70 °C) were collected in sterile deionized water and adjusted to a final concentration of 1×10^9^ CFU ml^-1^, which was equivalent to an optical density (OD_600_) of 1.0. The concentration was checked using a spectrophotometer (Hitachi U-1900, Tokyo, Japan).

### Testing of potato clones/cultivars for tuber soft rot resistance

Potato tubers were obtained from three environmental conditions by growing 176 clones of the mapping population, the parental clones P1 and P2, and two potato cultivars in the field for three consecutive years, 2017, 2018, and 2019. Tubers were stored after harvest in a regular potato storage house at 4-8 °C for approximately two (in 2019) or four months (in 2017, 2018). Tuber resistance to soft rot bacteria was tested in six independent experiments by the method previously described by Lebecka et al. (2018). In total, during the three years of testing, 36 tubers per individual (3 years × 2 dates × 2 replications × 3 tubers) and 72 tubers per parental clone and potato cultivar were tested. Tubers were rinsed in tap water and air-dried at room temperature 1 day before inoculation. Then, they were wounded with a steel rod (length 10 mm, Ø 2 mm) and inoculated with 10 μL of bacterial suspension, which was pipetted into a wound hole. The holes were sealed with Vaseline and a piece of parafilm. The tubers were externally sprayed with water, enclosed in boxes, and maintained in a climatic chamber for three days at 27 °C. The tubers were cut along the inoculation site, and the resistance was scored using two parameters: the disease incidence (DI), which was calculated as a proportion of tubers with rot symptoms, and the disease severity (DS), which was calculated as a mean weight of macerated tissue from tubers that showed symptoms of rotting.

### Testing of potato clones/cultivars for tuber starch content

Tuber starch content (TSC) was estimated through a comparison of the ratio of tuber weight in the air to tuber weight in water, according to Lunden (1956). The tests were conducted in two years, 2018 and 2019, with two technical replications. For each replicate, 6 tubers were weighed, and the same tubers were used in soft rot tests.

### Statistical analysis

Phenotypic data were collected for the two disease parameters DS and DI in three years and the TSC in two different years, and these parameters were analyzed as separate traits. Statistical analyses were performed using Statistica 10 software (Statsoft Inc. 2011). The normality of the distribution of the phenotypic data was tested by the Kolmogorov-Smirnov test. The effects of the potato clone, year, and interactions between these factors on each of three studied traits in the mapping population were estimated by two-factorial analysis of variance. For DS data, type IV analysis of variance was used, and for DI analysis, Bliss transformed data was used. A separate analysis of variance was performed for the data from parental clones and potato cultivars. The significance of differences among potato clones was estimated by Duncan’s test. The determination coefficients (*R^2^*) for tested traits were estimated from the sum of squares from analysis of variance. The skew of the phenotypic distributions of the tested traits was tested by the D’Agostino skewness test (Wessa 2017). To assess the reproducibility of test results between individual years and the correlation between the different traits, Pearson’s correlation coefficients were calculated. The broad-sense heritability coefficients (H_b_) of DS and TSC were estimated using the ANOVA results according to the following formula: H_b_ = σ2 g/(σ2 g + σ2 ge + σ2 e, σ2 g = (M1 – M2)/L; σ2 ge = M2— σe 2, where M1 = mean square of the effect of genotype, M2 = mean square of the effect of genotype × year interaction, L = number of years, σ2 e = mean square of the error (Domański et al. 2007).

### Genetic mapping and QTL analysis

Leaf samples of the progeny clones from the DS-13 mapping population, along with the parental clones P1 and P2, were collected from greenhouse-grown plants, quickly frozen in liquid nitrogen and ground with a micro pestle in Eppendorf tubes. DNA was isolated from 100 mg of powdered leaf tissue using a DNeasy Plant Mini Kit (Qiagen), with one modification in the Quick-Start Protocol: in steps 11 and 12 – 50 μL of AE buffer was used instead of 100 μL. Total DNA concentration and quality were estimated, and 20 μL samples of DNA adjusted to a concentration of 50 to 100 ng per μL were submitted to genotyping-by-sequencing (DArTseq^™^) at Diversity Arrays Technology Pty. Ltd. (Canberra, Australia) (http://www.diversityarrays.com/dart-application-dartseq). DArTseq^™^ genotyping was performed as described by Plich et al. (2018) and yielded 82 966 markers. The nucleotide sequences of the markers were located in the potato reference genome DM1-3 v4.04 (Hardigan et al. 2016) by a BLAST search (File S1). The resulting positions were encoded in the marker names by adding ‘chr’ followed by chromosome numbers from 01 to 12 and the positions (Mbp). Markers of presence and absence based on the variant (PAV) type, scored as 0 or 1, were used for map construction. Markers that i) did not segregate (showed a value of ‘1’ for less than 17 or more than 159 individuals), ii) had missing data for the parent(s) or a value of ‘0’ for both parents, iii) had missing data for 6 or more progeny individuals, iv) showed reproducibility < 1 were not included in the analyses. After import into JoinMap^®^ 4.1 (Van Ooijen 2012), markers that were identical in terms of segregation or were similar to each other (similarity ≥ 0.96) were also removed. Three PCR markers designed on the basis of candidate gene sequences (for metallocarboxypeptidase inhibitor, proteinase inhibitor type-2 CM7 K, and catechol oxidase) as described in Table S1 were scored in the mapping population and added to the DArTseq^™^ dataset (File S1). The reaction mixture using Thermo Scientific’s DreamTaq polymerase was as follows: 2 μl DreamTaq PCR buffer (Thermo Scientific), 1 μl 2 mM dNTPs, 0.4 μl (10 μM) F and R primers each, and 1 μl DNA template, supplemented with water to 20 μl. The reaction conditions were as follows: denaturation at 95 °C for 3 min; 35 amplification cycles of 95 °C for 30 s, 59 °C for 20 s and 72 °C for 2 min; and final synthesis at 72 °C for 10 min. PCR products of P01080 and M1BMR6 were subjected to restriction hydrolysis with the restriction enzyme Hinf I (Fermentas, Hanover). The enzymatic reaction was carried out at 37 °C for 3 h. Reaction mixture: 10 μL PCR product, 7.7 μL H_2_O, 2 μL buffer R, 0.3 μl Hinf I. The amplification products were separated in 1.4% agarose gels with Tris-borate-EDTA (TBE) buffer and ethidium bromide (0, 5 mg mL-1) and assessed under a UV trans illuminator. All three PCR products were sequenced to confirm their identity and size (Table S1) and were deposited in GenBank under the following accession numbers: MW509750, MW509751, MW509752. Linkage maps were calculated using a regression method with the following settings: CP (cross pollination/outbreeder full-sib family) population type, independence LOD (significance cut-off, LOD score > 3) as a grouping parameter and Haldane’s mapping function for the calculation of map distances. The DArTseq™ markers mapped to chromosomes other than those in the reference potato genome (PGSC DM1 −3 v 6.1) were removed manually from the maps. QTL analysis was performed as described previously (Śliwka et al. 2016, Hara-Skrzypiec et al. 2018) using interval mapping with MapQTL®6 software (Van Ooijen 2009). QTLs were detected using an LOD threshold ≥ 3.0 estimated from the cumulative distribution function of the maximum LOD on a chromosome for QTL analysis based on four QTL genotypes (van Ooijen 1999). In the DS-13 map, the average chromosome length was 97.5 cM, so the LOD threshold (α = 0.05) was between 2.9 (for a chromosome length of 50 cM) and 3.2 (100 cM).

## Data Availability Statement

The authors affirm that, all data necessary for confirming the conclusions of the article are present within the article, figures, tables, two additional supplemental files (File S1 contains DArT marker positions in potato reference genome DM1-3 v4.04, File S2 contains Supplement Genetic map and map summary), and two supplemental tables (Table S1 contains sequences of primers used in the genetic map and polymorphism of the candidate genes in parental forms of the mapping population DS-13, Table S2 contains correlation coefficients of disease severity, disease incidence and tuber starch content evaluated in three or two years of study of the mapping population) as well as three sequences of genes coding proteins: metallocarboxypeptidase inhibitor, proteinase inhibitor type-2 CM7 K, and catechol oxidase in the NCBI GenBank under the accession numbers: MW509750, MW509751, MW509752.

## Results

### Evaluation of DS, DI and TSC

Potato tubers were wound inoculated with *D. solani*, and three days post inoculation, they showed two types of reactions: symptoms of wet macerated tissue or dry, necrotic and suberized lesions (Fig. 1). The distributions of the phenotypic parameters DS, DI and TSC in the DS-13 segregating population are presented in Fig. 2. The mean DS of the progeny genotypes was distributed normally and positively skewed according to the D’Agostino skewness test. The phenotypic distribution of DI deviated significantly from normality according to the Kolmogorov-Smirnov test. The results of two-way ANOVA indicated the significant effect of genotype, year, and the genotype by year interaction on the DS, DI and TSC values (all p values < 0.001) (Table 1). The broad sense heritability coefficients were 0.50 for DS, 0.24 for DI and 0.79 for TSC (Table 1). The descriptive statistics of DS, DI and TSC evaluated in the parental clones, their F1 progeny in population DS-13, and two potato cultivars are presented in Table 2. P1 expressed a significantly higher level of resistance to inoculation with *D. solani* than P2 and the moderately resistant standard cultivar Glada, as indicated by Duncan’s post hoc test. The susceptible parent P2 did not differ from the highly susceptible standard cultivar Irys in terms of *D. solani* resistance. The Pearson’s correlation coefficients between parameters/traits and years are presented in Table S2. The scoring of the progeny individuals was more reproducible between the two seasons for the TSC than for DS and DI, tested in three seasons. The correlation coefficients of the year means ranged from 0.48 to 0.69 (p<0.001) for DS, from 0.33 to 0.46 (p<0.001) for DI and 0.81 (p<0.001) for the TSC. The correlation coefficients between mean DS and mean DI were 0.53 (p<0.001), between DS and TSC were −0.26 (p<0.001), and between DI and TSC were −0.19 (p<0.01).

**Fig. 1.**
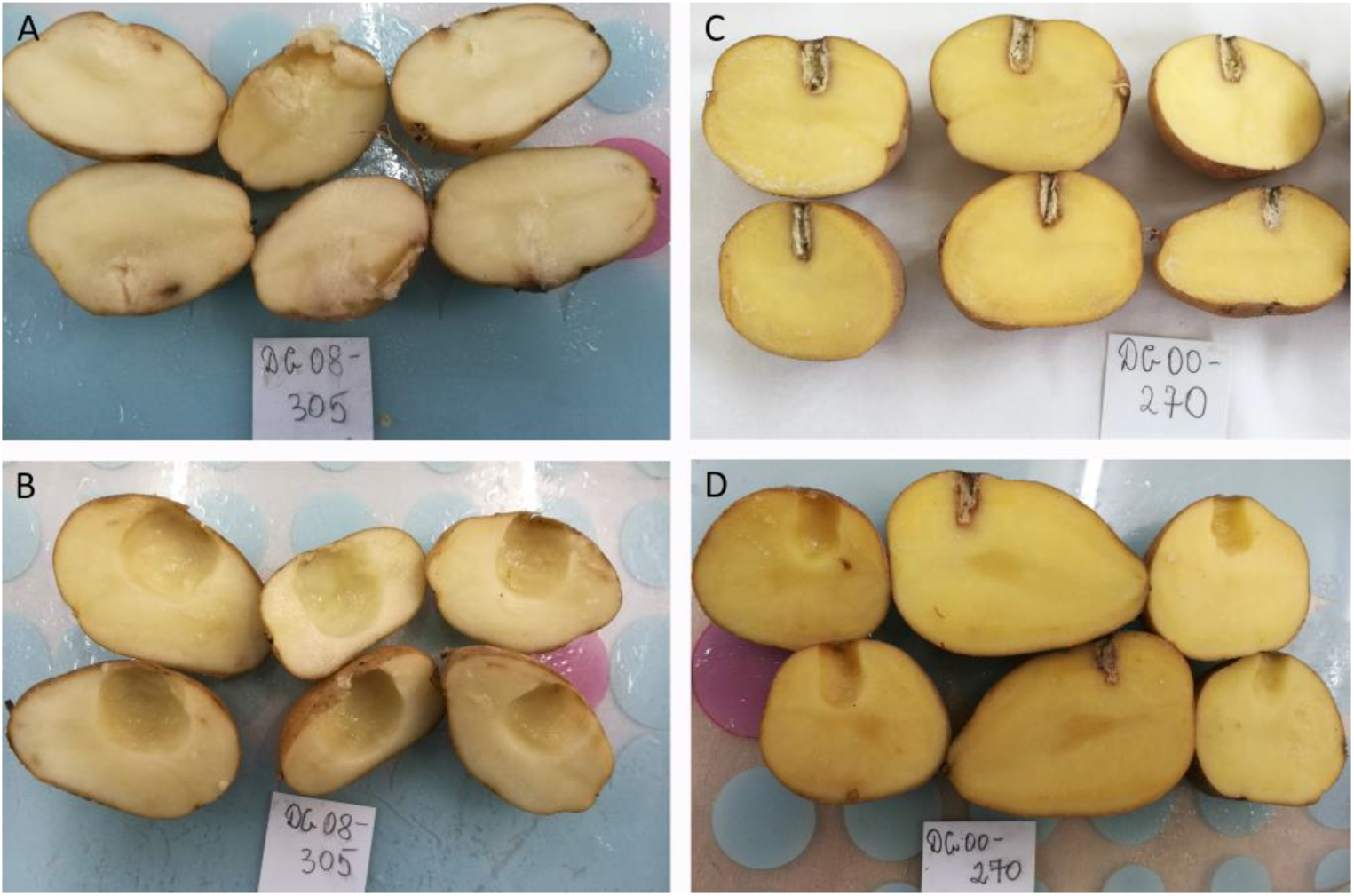
Potato tubers showing a range of responses to wound inoculation with *Dickeya solani*. At 72 h post inoculation, the tubers were cut in half, macerated tissue was removed, and two halves of the same tuber are shown: a) the maternal parent of the mapping population P1 shows a high level of resistance expressed as wound-healing resistance (two upper tubers cut in half) and as a low weight of rotten tissues (the third one cut in half), b) the pollen parent P2 shows a susceptible reaction expressed as rotten tissue (removed from tubers), c) the F1 progeny, the diploid clone DS-230, expressed high resistance, d) the F1 progeny, the diploid clone DS-186, was susceptible to *D. solani*.

**Fig. 2.**
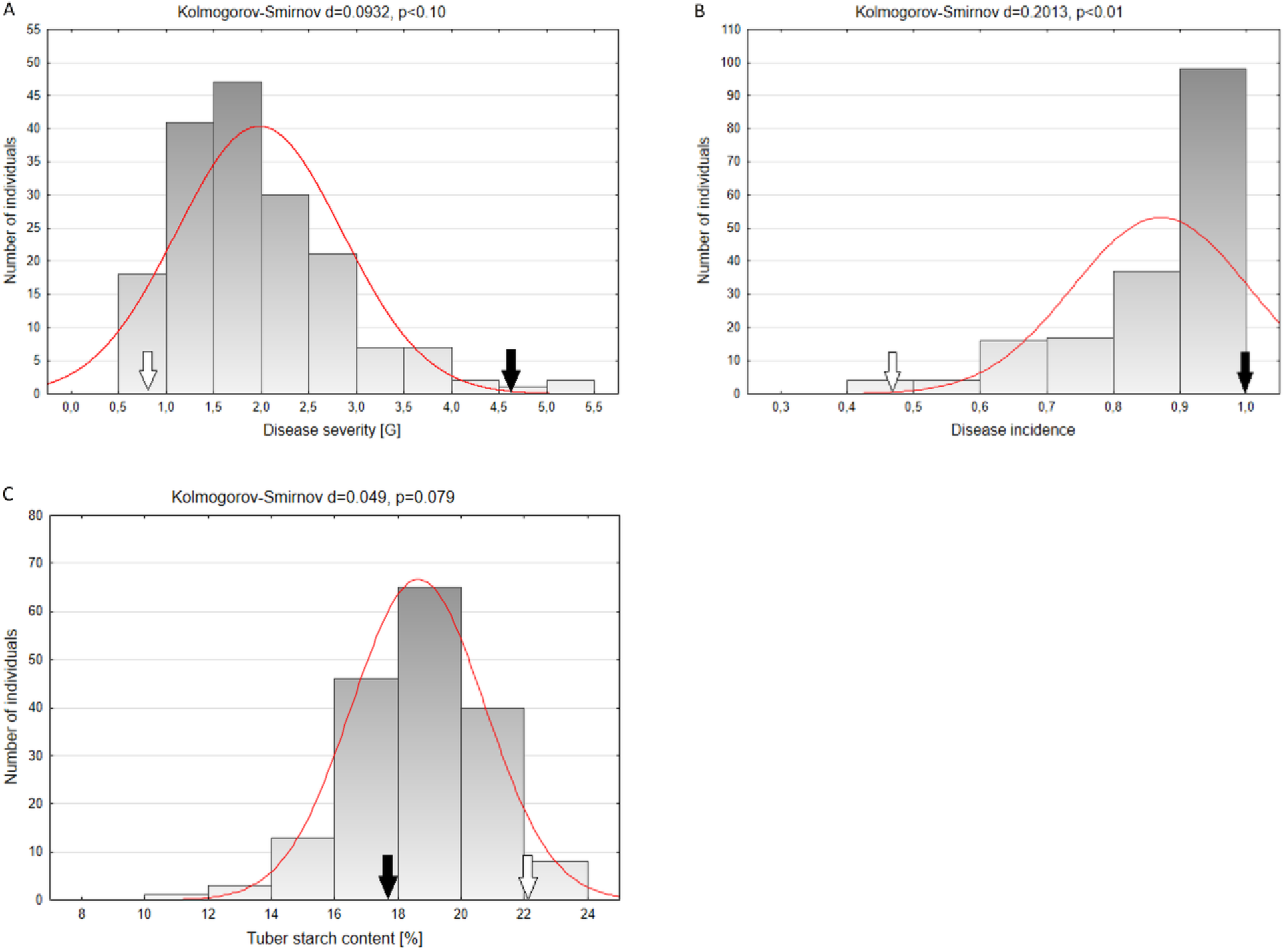
Histograms of frequency distributions of A) average disease severity expressed as the weight of rotted tuber tissue, B) average disease incidence expressed as the proportion of tubers with symptoms of rot in the mapping population consisting of 176 individuals, obtained during six independently repeated soft rot resistance evaluation experiments performed in three consecutive years, 2017, 2018 and 2019, C) average tuber starch content in the mapping population, evaluated in two experiments performed in two years, 2018 and 2019. White and black arrows indicate the three- or two-year mean values of the P1 and P2 clones, respectively. The results of normality tests are shown above the charts; fit to the normal curve is indicated by a line.

### QTL analysis

After a quality check, 7056 markers segregating in population DS-13 were imported into JoinMap^®^ 4.1. A BLAST search revealed the localization of 5156 (73%) markers in the potato reference genome DM1-3 v4.04 (File S1). A fraction of markers (4.3%) was located among the unanchored sequences of the reference genome (Chr00 or ChrUn). After preliminary linkage analyses, the markers that were linked to chromosomes other than those in the reference genome were also deleted. The final DS-13 genetic map of both parents was 1169.5 cM in total length and consisted of 2231 DArTseq^™^ and six PCR markers (File S2). The average number of markers per chromosome was 186.4 and ranged from 134 on chromosome X to 276 on chromosome II. The average chromosome length was 97.5 cM, and it varied from 79.3 cM (chromosome V) to 123.3 cM (chromosome I) (File S2).

The DS-13 genetic map and the phenotyping results were used for QTL interval mapping with MapQTL^®^6 software. Strong and reproducible QTLs for soft rot resistance were detected on chromosomes II and IV (Table 3). The QTL on chromosome II affected DS only and was significant in three out of four datasets (DS2017, DS2019 and DS17-19). The mean DS17-19 dataset spanned the 8.0-33.1 cM region and explained up to 16.5% of the variance (LOD 6.87). The QTL on chromosome IV was significant for both DS and DI in all datasets from particular years and in the mean 17-19 datasets. Its effect explained 22.4% and 22.9% of the variance in DS and DI, respectively and was significant for both parameters in a large portion of chromosome IV (DS17-19: 0.0-69.1 cM; DI17-19: 0.0-83.6 cM; Fig. 3). The LOD charts for DS and DI in particular datasets overlapped and peaked between 40 and 55.2 cM, with the exceptions of DS2019 and DI2017 (Fig. 3, Table 3).

**Fig. 3.**
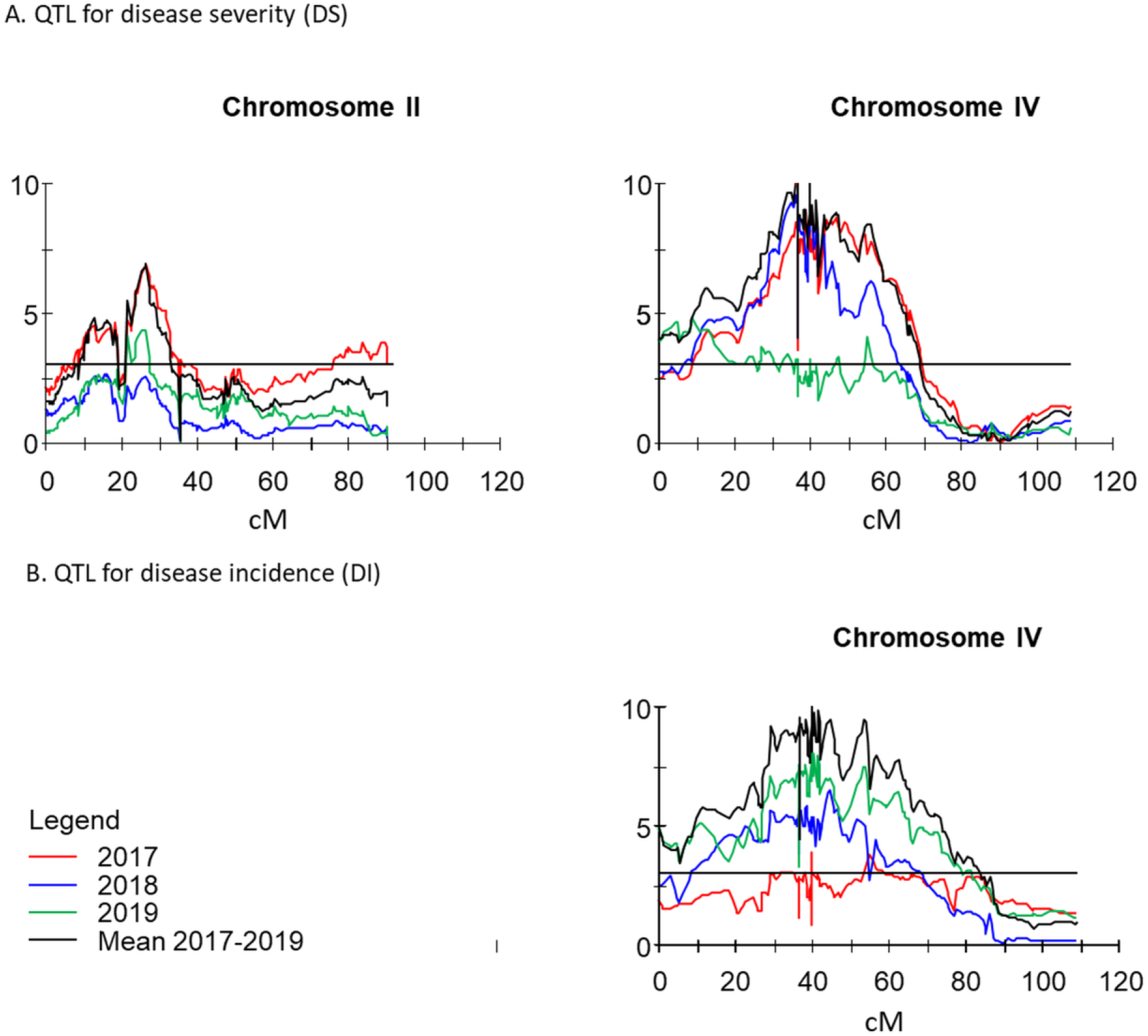
The most important quantitative trait loci for potato tuber resistance to soft rot caused by *Dickeya solani* in the diploid potato mapping population DS-13: A) mean disease severity (2017-2019), and B) mean disease incidence (2017-2019). LOD significance thresholds for QTL analysis are indicated by a horizontal line (LOD = 3.0).

Additionally, minor QTLs for soft rot resistance were detected in only one or two of the four datasets, and their effects explained no more than 10.4% of the variance (Table 3). Minor QTLs for DS were found on chromosomes II (76.0-90.4 cM), V and X, while for DI, minor QTLs were located on chromosomes I and X in regions not overlapping those of the QTLs for DS (Table 3).

Three sequence-specific markers derived from genes of proteins that were differentially expressed between the resistant P1 and susceptible P2 parental clones identified by Lebecka et al. (2019) at the early stage of *D. solani* infection were mapped in the DS-13 population. Their genetic positions are given in Table S1. None of the markers were mapped within QTLs for DS or DI.

Ten significant QTLs for the TSC were mapped to eight potato chromosomes. The most replicable QTLs were located on chromosomes III, XI, and XII, and two QTLs were located on chromosome V (Table 4). Only one QTL was mapped to chromosome IV (significant in 2018), but it was located in a different region than the QTLs for DS and DI.

## Discussion

There is no standardized method for the evaluation of potato tuber soft rot resistance after artificial inoculation with bacteria. The methods differ in terms of inoculation technique, incubation conditions and evaluation method, as described by Lebecka (2017). A point inoculation method including wound inoculation of whole potato tubers by using micropipette tips was proposed by Priou et al. (1992) as a standard method; however, the results of the ring test performed by different European laboratories differed among laboratories (Allefs et al. 1993). In our research, we used a point inoculation method described by Laurila et al. (2008) with modifications (Lebecka et al. 2018). Most researchers assess the resistance of potato tubers using one parameter, the size of rotten tissue (DS in our study), which corresponds to the resistance of the tuber tissue to maceration by bacteria. An additional resistance scoring criterion was first applied by Chung et al. (2017), wherein in addition to lesion size, lesion type was scored, where a score of 1 indicates a lesion blocked from expansion and corresponds to a barrier-like resistance mechanism - a rapid form of wound healing. The DI parameter used in this study indicates the frequency of tubers with symptoms of infection corresponding to this type of resistance (Lebecka et al. 2019). We used these two parameters because in our experiments, in addition to most tested tubers showing symptoms of soft rot, there were also tubers that exhibited a quick wound-healing response at the wound area, resulting in suberized lesions that successfully protected the tuber against pectinolytic bacterial activity. Suberization is induced by wounds in potato tubers as a part of the overall wound-healing process to cover the damaged areas. The final form of this healing layer, the suberin phenolic domain, provides resistance to infection by soft rot bacteria at the wound site (Lulai and Corsini, 1998). The clone P1 used in this study and the diploid wild potato relative M6 expressed a barrier-like resistance mechanism to soft rot bacteria (Chung et al. 2017). The process of internal suberization of tuber parenchyma cells can be induced by a pathogen without a wound signal, which has been shown in an experiment with *Verticillium dahliae*, which infects tubers through xylem vessels (Lulai et al. 2005). Therefore, it might be difficult to separate the effect of wounding from the effect of bacterial infection on this type of barrier-like resistance.

Resistance to soft rot is highly heritable in clones representing the 2^nd^ source of resistance, described by Lebecka and Zimnoch-Guzowska (2004), with a high coefficient of broad-sense heritability (0.92), suggesting the simple manner of the inheritance of soft rot resistance. In this study, the broad-sense heritability of DS was moderate (*H^2^* = 0.50) and similar to the heritability of resistance to *Erwinia chrysanthemi* (*Dickeya* spp. in present) estimated in diploid populations of *S. stenotomum* and *S. phureja* (*H^2^* = 0.59) by using the diameter of the rotten area as a measure of the degree of resistance (Wolters and Collins 1991). The broad-sense heritability of DS was also similar or lower than the heritability for resistance to common scab, other bacterial disease caused by *Streptomyces scabies*, evaluated in field trials in a diploid population of *S. tuberosum* and *S. chacoense* (*H^2^* in range from 0.48 to 0.72) (Braun et al. 2017), and lower than that estimated for common scab resistance in potato clones and breeding lines (*H^2^* = 0.81) (Yuan et al. 2020). The broad-sense heritability of DI in our study was low (*H^2^* = 0.24). The results confirmed a large role of the environment in soft rot resistance; nevertheless, we were able to consistently select highly resistant and highly susceptible individuals in the mapping population for genomic studies. In addition, the P1 parent is a good candidate for use as a parent in 2*x* – 4*x* crosses due to its ability to produce large pollen grains. When 2*n* gametes are formed during the First Division Restitution, they transfer both additive and nonadditive parts of the genetic variance to the tetraploid progeny (Mendiburu and Peloquin 1977).

To describe the phenotypic reaction of potato tubers wound-inoculated with bacteria, two different parameters were used: DS and DI. QTL analysis of the DS-13 mapping population revealed prominent QTLs for DS on potato chromosomes II and IV and one QTL for DI on chromosome IV that was mapped to an overlapping position with the QTL for DS. In total, we detected eight QTLs for soft rot resistance caused by *D. solani* on five potato chromosomes. Overlapping QTLs for different parameters can result either from pleiotropic effects of a single gene or from the effects of closely linked but otherwise unrelated genes (Gebhardt 2005). The phenotypic correlation between DS and DI was positive and moderate (Pearson’s *r* = 0.53, p<0.001). Our assumption that DS and DI are only partly related is supported by the finding of some QTLs for both parameters in the overlapping position on chromosome IV and some unique QTLs for each parameter. We recommend using both parameters for more accurate characterization of potato reactions to soft rot bacteria.

Precise comparison of the QTL position between our work and the first mapping study of resistance to soft rot caused by *P. atrosepticum* (Zimnoch-Guzowska et al. 2000) is not possible due to the use of different marker systems and lack of consensus maps. Zimnoch-Guzowska et al. (2000) revealed the complexity of the trait: 13 putative QTLs were identified on 10 linkage groups. None of these QTLs were detected in any of the datasets in this study. The strongest and most reproducible QTL was mapped to chromosome I and explained up to 19% of phenotypic variance. Among others, QTLs on chromosomes II and IV were identified and explained up to 5.5 and 8.6% of the variance, respectively (Zimnoch-Guzowska et al. 2000), and these QTLs may correspond to those described in our work, since we hypothesized that resistance against different pectinolytic bacteria may be at least partially based on common mechanisms. The smaller number of QTLs in our study than in the work by Zimnoch-Guzowska et al. (2000) and the differences in the QTL landscape are likely caused by the use of genetically diversified plant material and different pathogen species and phenotyping methods.

Resistance QTLs for other bacterial diseases of potato, such as common scab, have recently been mapped in a genome–wide association study in 143 Canadian potato cultivars and advanced breeding clones (Yuan et al. 2020). Three QTLs for potato tuber resistance to this pathogen, explaining 21%, 8%, and 12% of phenotypic variance, were identified on potato chromosomes II, IV and XII, respectively. The QTL on chromosome II spanned a region corresponding to 36.5-36.7 Mbp of chr02 in the reference genome (Yuan et al. 2020), while our major QTL for DS corresponded to the 21.7-34.4 Mbp region. Although the peak of our second major QTL for DS and DI on chromosome IV was located at a position ascribed to approximately 50.2 Mbp in the reference genome, the effect of this QTL also extended to the region described by Yuan et al. (2000) as affecting common scab resistance (chr04: 11.6-12.6 Mbp), indicating that this region of the potato genome may be generally involved in resistance to bacterial pathogens.

The infection of potato tubers in favorable conditions is rapid; the duration of the asymptomatic phase in potato tubers infected with *D. solani* is 8 h (Lebecka et al. 2019), and in chicory leaves infected by *D. dadantii* 3937, it was estimated to be between 8 and 12 h (Effantin et al. 2011). In our recent paper, we detected 9 proteins with significant fold changes between DG 00-270 (P1 in this study) and DG 08-305 (P2) 8 h post inoculation with *D. solani* (Lebecka et al. 2019). Here, for three genes that encode the selected proteins, metallocarboxypeptidase inhibitor, proteinase inhibitor type-2 CM7 K, and catechol oxidase, DNA markers were developed and mapped in population DS-13. None of the genetic factors appeared to be colocalized with trait QTLs. Protein abundance is controlled by variation at the coding gene itself and by variation mapping to other regions of the genome (Foss et al. 2007). Comparative analyses of phenotypic QTLs with protein QTLs can provide a more direct link between protein abundance and traits.

The reproducibility of the analyzed TSC over two years was high (Pearson *r* = 0.81, *p*<0.05) and more stable over years than that observed for DS and DI. The same potato tubers that were evaluated for TSC in 2018 and 2019 were evaluated for DS and DI. At the phenotypic level, the correlation coefficients between TSC and DS or DI in each of the years were significant but low. The mean correlation coefficient between TSC and DS (Pearson’s *r* = 0.23, *p*<0.05) was two times lower than the correlation coefficient between dry matter and lesion diameter score obtained in the diploid mapping population of *S. tuberosum* clones and *S. chacoense* clones in the studies of Chung et al. (2017). The low correlation obtained in this study is consistent with the fact that QTLs for TSC were located on different chromosomes than QTLs for soft rot resistance. Earlier reports of the presence of potato cultivars with a high content of dry matter together with lower susceptibility to soft rot might be explained by the effect of previous selection of genetically unlinked traits. It may also be possible that there is an indirect relationship between these traits due to the influence of starch content on the physical characteristics and chemical properties of potato tubers.

When investigating the metabolite profile of extract from healed potato tuber wounds, time-dependent changes in biomarker structure and distinctive patterns for different potato cultivars were revealed, with the highest antibacterial activity against *P. carotovorum* observed for the polar extracts from day 0 post wounding (6 h post wounding, personal communication, Dastmalchi) (Dastmalchi et al. 2019). Decreasing antimicrobial activity occurs 3-7 days post wounding, prior to the formation of a physical periderm barrier. We hypothesized that P1, a source of high resistance to pectinolytic bacteria that is higher than the resistance level found in potato cultivars, might have different metabolic profiles in the early phase of infection, leading to a quick wound-healing response and slower rot of tuber tissue than that observed in potato cultivars. Genomic approaches will be undertaken in future studies to verify this hypothesis and to select and validate candidate genes located in QTL regions that were identified in this study.

We described a new source of high resistance to *D. solani* originating from wild and primitive *Solanum* species, QTLs conferring two types of resistance, quick wound healing (measured as low DI) and slow rot of tuber tissue (DS). The high level of resistance in P1 and its resistant progeny, exhibited under disease-favorable conditions, should be manifested as an even higher level of resistance under less suitable conditions for disease development during standard potato tuber storage. The lack of a linkage between the TSC and DS or DI enables improvement of resistance and maintenance of high starch contents in potato breeding programs.

## Declarations

### Ethics approval

Not applicable

### Consent to participate

Not applicable

### Consent for publication

Consent for publication has been obtained from all authors

### Code availability

Not applicable

## Author Contributions Statement

RL conceived and coordinated the project, selected parental lines, performed crossing, conducted phenotyping, performed part of the statistical analysis, and co-wrote the manuscript; JŚ constructed the genetic map, conducted the QTL analysis and co-wrote the manuscript; AGU performed marker screening; KS performed DNA extraction and sequencing of PCR markers; WM contributed to manuscript writing. All authors contributed to manuscript revision and read and approved the submitted version.

## Abbreviations

ANOVA: Analysis of variance
BLAST: Basic local alignment search tool
CFU: Colony forming units
QTL: Quantitative trait loci
CP: Cross pollination
DArTseq: Diversity array technology
DI: Disease incidence
DS: Disease severity
H_b_: Broad-sense heritability coefficient
LOD: Logarithm of the odds
OD: Optical density
PAV: Presence and absence variant
PGSC: Potato genome sequencing consortium
qRT-PCR: Real-time quantitative polymerase chain reaction
TBE: Tris-borate-ethylenediaminetetraacetic acid buffer
TSC: Tuber starch content

## Acknowledgements

This research was financially supported by a grant from the Ministry of Agriculture and Rural Development, Poland (HOR.hn.802.19.2018 task number 56). We would like to express our gratitude to Prof. Ewa Zimnoch-Guzowska (Plant Breeding and Acclimatization Institute) for her valuable comments on the manuscript and her editorial work.

## References

Bethke PC, Halterman DA, Jansky SH (2019) Potato germplasm enhancement enters the genomics era. Agronomy 9:575 https://doi.org/10.3390/agronomy9100575

Biehn WL, Sands DC, Hankin L (1972) Relationship between percent dry matter content of potato tubers and susceptibility to bacterial soft rot. Phytopathol 62: 747 (Abstract)

Braun SR, Endelman JB, Haynes KG, Jansky SH (2017) Quantitative trait loci for resistance to common scab and cold-induced sweetening in diploid potato. Plant Genome 10(3) https://doi.org/10.3835/plantgenome2016.10.0110

Chung YS, Kim C, Jansky S (2017) New source of bacterial soft rot resistance in wild potato (*Solanum chacoense*) tubers. Genetic Res Crop Evol 64(8): 1963–1969 https://doi.org/10.1007/s10722-017-0487-3

Charkowski A, Sharma K, Parker ML, Secor GA, Elphinstone J (2020) Bacterial Diseases of Potato. The Potato Crop. Springer International Publishing 2020:351–88. Available from: https://doi.org//10.1007/978-3-030-28683-5_10

Chung YS, Kim C, Jansky S. (2017) New source of bacterial soft rot resistance in wild potato (*Solanum chacoense*) tubers. Genet Resour Crop Evol 64:1963–1969 https://doi.org/10.1007/s10722-017-0487-3

Czajkowski R, Grabe GJ, Van der Wolf JM (2009) Distribution of *Dickeya* spp. and *Pectobacterium carotovorum* subsp. *carotovorum* in naturally infected seed potatoes. Eur J Plant Pathol 125:263–275

Dastmalchi K, Perez-Rodriguez M, Lin J, Barney Y, Stark RE (2019) Temporal resistance of potato tubers: antibacterial assays and metabolite profiling of wound healing tissue extracts from contrasting cultivars. Phytochemistry 159:75–89

Domański L, Michalak K, Zimnoch-Guzowska E (2007) Variation of blackspot susceptibility of the selected potato cultivars. Biuletyn IHAR 246:145–149 http://biblioteka.ihar.edu.pl/biuletyn_ihar.php?field[slowa_kluczowe]=&field[autor]=&id=40&idd=884&podzial_id=2&podzial_idd=#lib

Effantin G, Rivasseau C, Gromova M, Bligny R, Hugouvieux-Cotte-Pattat N (2011) Massive production of butanediol during plant infection by phytopathogenic bacteria of the genera *Dickeya* and *Pectobacterium*. Mol Microbiol 82:988–997

Foss EJ, Radulovic D, Shaffer SA, Ruderfer DM, Bedalov A, Goodlett DR, Kruglyak L (2007) Genetic basis of proteome variation in yeast. Nat Genet 39(11): 1369–1375 https://doi.org/10.1038/ng.2007.22

Gebhardt C (2005) Potato genetics: molecular maps and more. In: Lörz H, Wenzel G (eds) Molecular marker systems in plant breeding and crop improvement. Springer, Berlin/Heidelberg, pp 215–227

Golanowska M, Galardini M, Bazzicalupo M, Hugouvieux-Cotte-Pattat N, Mengoni A, Potrykus M, Slawiak M, Lojkowska E (2015) Draft genome sequence of a highly virulent strain of the plant pathogen *Dickeya solani*, IFB0099. Genome Announc 3. https://doi.org/10.1128/genomeA.00109-15

Hara Skrzypiec A, Śliwka J, Jakuczun H, Zimnoch-Guzowska E (2018) Quantitative trait loci for tuber blackspot bruise and enzymatic discoloration susceptibility in diploid potato. Mol Genet Genomics 293(2): 331–342

Hardigan MA, Crisovan E, Hamilton JP, Kim J, Laimbeer P, Leisner CP, Manrique-Carpintero NC, Newton L, Pham GM, Vaillancourt B, Yang X, Zeng Z, Douches DS, Jiang J, Veilleux RE, Buell CR (2016) Genome reduction uncovers a large dispensable genome and adaptive role for copy number variation in asexually propagated *Solanum tuberosum*. Plant Cell 28(2):388–405

Hugouvieux-Cotte-Pattat N (2016) Metabolism and virulence strategies in *Dickeya*-host interactions. Prog Mol Biol Transl Sci 142:93–129 https://doi.org/10.1016/bs.pmbts.2016.05.006

Kraepiel Y, Barny MA (2016) Gram-negative phytopathogenic bacteria, all hemibiotrophs after all? Mol Plant Pathol 17:313–316

Lebecka R, Michalak K (2017) Reakcja bulw wybranych odmian ziemniaka na porażenie przez wysokowirulentny szczep bakterii *Dickeya solani*. (in Polish) Ziemniak Polski 3: 18–23 http://biblioteka.ihar.edu.pl/ziemniak_polski.php?field[slowa_kluczowel=&field[autorl=&id=60&idd=498&podzialid=&podzialidd=

Lebecka R, Zimnoch-Guzowska E (2004) The inheritance of resistance to soft rot (*Erwinia carotovora* subsp. *atroseptica*) in diploid potato families. Amer J Potato Res 81:395–401

Lebecka R, Zimnoch-Guzowska E, Kaczmarek Z, (2004) Resistance to soft rot (*Erwinia carotovora* subsp. *atroseptica*) in tetraploid potato families obtained from 4*x*-2*x* crosses. Amer J Potato Res 82:203–210

Lebecka R, Flis B, Murawska Z (2018) Comparison of temperature effects on the in vitro growth and disease development in potato tubers inoculated with bacteria *Pectobacterium atrosepticum*, *P. *carotovorum* subsp. *carotovorum* and *Dickeya solani**. J Phytopathol 166:654–662

Lebecka R, Kistowski M, Dębski J, Szajko K, Murawska Z, Marczewski W (2019) Quantitative proteomic analysis of differentially expressed proteins in tubers of potato plants differing in resistance to *Dickeya solani*. Plant and Soil 441: 317–329 https://doi.org/10.1007/s11104-019-04125-7

Lulai EC (2005) Non-wound-induced suberization of tuber parenchyma cells: a physiological response to the wilt disease pathogen *Verticillium dahliae*. Amer J Potato Res 82:433–40 https://doi.org/10.1007/BF02872221

Lulai EC and Neubauer JD (2014) Wound-induced suberization genes are differentially expressed, spatially and temporally, during closing layer and wound periderm formation. Postharvest Biol Tec 90:24–33

Lulai EC, Campbell LG, Fugate KK, McCue KF (2016) Biological differences that distinguish the 2 major stages of wound healing in potato tubers. Plant Signal Behav 11(12) p.e1256531

Lulai EC and Corsini DL (1998) Differential deposition of suberin phenolic and aliphatic domains and their roles in resistance to infection during potato tuber (*Solanum tuberosum* L.) wound-healing. Physiol Mol Plant Pathol 53:209–222

Lunden PA (1956) Underldokerd over forholder mellom popetens spesifikka vekt og deres torvstoff og Stivelsesinhold Forhl. Forsok Landbruket 7:81–107

Ma B, Hibbing ME, Kim HS, Reedy RM, Yedidia I, Breuer J, Breuer J, Glasner JD, Perna NT, Kelman A, Charkowski AO (2007) Host range and molecular phylogenies of the soft rot enterobacterial genera pectobacterium and dickeya. Phytopathol 2007 97(9):1150–63 https://doi.org/10.1094/PHYTO-97-9-1150

Mansfield J, Genin S, Magori S, Citovsky V, Sriariyanum M, Ronald P, Dow M, Verdier V, Beer SV, Machado MA, Toth I, Salmond G, Foster GD (2012) Top 10 plant pathogenic bacteria in molecular plant pathology. Molec Plant Pathol 13:614–629

Mendiburu AO, Peloquin SJ (1977) The significance of 2N gametes in potato breeding. Theor Appl Genet 49:53–61

Pham GM, Hamilton JP, Wood JC, Burke JT, Zhao H, Vaillancourt B, Ou S, Jiang J, Buell CR (2020) Construction of a chromosome-scale long-read reference genome assembly for potato, GigaScience 9, 9 giaa100, https://doi.org/10.1093/gigascience/giaa100

Plich J, Przetakiewicz J, Śliwka J, Flis B, Wasilewicz-Flis I, Jakuczun H, Zimnoch-Guzowska E (2018) Novel gene *Sen2* conferring broad-spectrum resistance to *Synchytrium endobioticum* mapped to potato chromosome XI. Theoret Appl Genet 131(11): 2321–2331

Slawiak M, Łojkowska E, Van der Wolf JM (2009) First report of bacterial soft rot on potato caused by *Dickeya* sp. (syn. *Erwinia chrysanthemi*) in Poland. Plant Pathol 58:794

Śliwka J, Sołtys-Kalina D, Szajko K, Wasilewicz-Flis I, Strzelczyk-Żyta D, Zimnoch-Guzowska E, Jakuczun H, Marczewski W (2016) Mapping of quantitative trait loci for tuber starch and leaf sucrose contents in diploid potato. Theor Appl Genet 121: 131–140

Toth IK, van der Wolf JM, Saddler G, Lojkowska E, Hélias V, Pirhonen M, Tsror L, Elphinstone JG (2011) *Dickeya* species: an emerging problem for potato production in Europe. Plant Pathol 60:385–399

Tzeng KC, McGuire RG, Kelman A (1990) Resistance of tubers from different potato cultivars to soft rot caused by *Erwinia carotovora* subsp. *atroseptica*. Amer J Potato Res 67:287–305

Van Ooijen JW (2012) JoinMap ® 4.1. Software for the calculation of the genetic linkage maps in experimental populations of diploid species. Kyazma BV. Wageningen

Van Ooijen JW (2009) MapQTL ® 6. Software for mapping of quantitative trait loci in experimental populations of diploid species. Kyazma BV. Wageningen

Van Ooijen JW (1999) LOD significance thresholds for QTL analysis in experimental populations of diploid species. Heredity 83:613–624

Wessa P (2017) Skewness and Kurtosis Test (v1.0.4) in Free Statistics Software (v1.2.1), Office for Research Development and Education https://www.wessa.net/rwasp_skewness_kurtosis.wasp

Wolters PJ, Collins WW (1991) Inheritance of *Erwinia* resistance and the correlation between resistance and specific gravity in diploid potatoes. In: Abstracts of papers presented at the 75th Annual Meeting of The Potato Association of America Spokane, Washington State August 11-15, 1991. Amer Potato J 68:595–643 https://doi.org/10.1007/BF02853713

Wright PJ, Triggs CM, Anderson JAD (2005) Effects of specific gravity and cultivar on susceptibility of potato (*Solanum tuberosum*) tubers to blackspot bruising and bacterial soft rot. New Zeal J Crop Hort, 33:4, 353–361, DOI:10.1080/01140671.2005.9514370

Yuan J, Bizimungu B, De Koeyer D, Rosyara U, Wen Z, Lagüe M (2020) Genome-wide association study of resistance to potato common scab. Potato Res 63:253–266

Zimnoch-Guzowska E, Łojkowska E (1993) Resistance to *Erwinia* in diploid potatoes with high starch content. Potato Res 36: 177–182

Zimnoch-Guzowska E, Marczewski W, Lebecka R, Flis B, Schaefer-Pregl R, Gebhardt C (2000) QTL analysis of new sources of resistance to *Erwinia carotovora* ssp. *atroseptica* in potato done by ALFP, RFLP, and resistance-gene-like markers. Crop Sci 40:1156–1167

